# Annexin A1 as a key modulator of lung inflammation during coronavirus infections

**DOI:** 10.1101/2025.02.12.637941

**Authors:** Filipe Resende, Celso Martins Queiroz-Junior, Fernando Roque Ascenção, Ian de Meira Chaves, Larisse de Souza Barbosa Lacerda, Felipe Rocha, Danielle Teixeira, Izabela Galvão, Victor Costa, Talita Fonseca, Arthur Gualberto, Ana Luiza de Castro Santos, Jenniffer Martins, Erick Bryan de Sousa Lima, Adelson Héric Alves Monteiro, Isabella Zaidan, Laís C. Grossi, Pedro Augusto Carvalho Costa, Vinicius Amorim Beltrami, Lirlândia P. Sousa, Pedro Pires Goulart Guimarães, Gabriel Campolina-Silva, Mauro M Teixeira, Vanessa Pinho, Vivian V Costa

## Abstract

Exacerbated inflammation is a major contributor to tissue damage and mortality in infectious diseases, including SARS-CoV-2. The resolution phase of inflammation is critical for restoring tissue homeostasis following an injury. Annexin A1 (AnxA1) is a ubiquitous protein that plays a fundamental role in the resolution of inflammation, including in preclinical models of infectious disease. Here, we investigated the role of AnxA1 in coronavirus infection and its potential as a host-targeted therapeutic strategy against SARS-CoV-2. Wild-type (WT) and AnxA1 knockout (AnxA1KO) mice were intranasally infected with the murine betacoronavirus MHV-3 to study the endogenous role of AnxA1. Immunohistochemistry and Western blot analyses in the lungs of MHV-3-infected mice revealed increased AnxA1 expression and its cleavage, which was associated with neutrophilic infiltration (Ly6G+ cells) mainly in peribronchiolar and perivascular regions. AnxA1-deficient mice exhibited higher neutrophilic infiltration and lung damage, alongside increased CXCL1 production in the lungs, when compared to WT-infected mice. In a murine model of SARS-CoV-2 infection in K18-hACE2 mice, we found increased AnxA1 cleavage associated with lung inflammation. Treatment of SARS-CoV-2-infected K18-hACE2 mice with the AnxA1-mimetic peptide, Ac_2-26_, reduced lung damage and lethality, without altering the host ability to deal with viral replication. Notably, Ac_2-26_-treated mice exhibited similar levels of protection to that afforded by the nucleotide analogue Remdesivir, following SARS-CoV-2 infection. Our findings highlight the protective role of the endogenous AnxA1 in mitigating coronavirus-induced lung inflammation and underscore the therapeutic potential of AnxA1 mimetic Ac_2-26_ as a host-targeted therapy against SARS-CoV-2.

## Introduction

Betacoronaviruses including MERS-CoV, SARS-CoV, and SARS-CoV-2, are characterized by high infection and mortality rates, posing a significant threat to global public health. COVID-19 (Coronavirus Disease 2019) has particularly underscored the risks associated with emerging and re-emerging viral diseases. These viruses induce dysregulated inflammation, which contributes to tissue damage and elevated mortality rates, exemplifying a recurring issue in severe infections (Mackay and Arden, 2015; Chan-Yeung and Xu, 2003; Msemburi et al., 2023). As a result, the idea of therapies targeting exacerbated inflammation have emerged as a compelling host-target alternative for treating infectious diseases, such as the use of dexamethasone in severe COVID-19 (Garcia et al., 2010; Sousa et al., 2020; Maskin et al., 2022).

Excessive inflammatory response and consequent disease can be due to failure of resolutive circuits of inflammation. The resolution of inflammation unfolds after early phases of inflammatory response, in which leukocytes undergo a shift in their phenotype driven by pro-resolving mediators to restore homeostasis. These endogenous mediators include lipids, such as Lipoxin A4 and Resolvins, and proteins such as Annexin A1 (AnxA1) (Perretti et al., 2015; Sugimoto et al., 2019).

AnxA1 is a 346-amino acid protein ubiquitously expressed in mammals that exerts its resolutive effects mainly by binding to Formyl Peptide Receptor 2 (FPR2), a G-protein coupled receptor expressed by diverse leukocytes (Sugimoto et al., 2016). Notably, in addition to its role in inflammation resolution, AnxA1 also shows anti-inflammatory effects. Using genetic ablated mice for AnxA1 and AnxA1-mimetic peptide (Ac_2-26_), our research group and others have shown the importance of this protein in controlling inflammation and promoting the resolution phase across various preclinical models, such as bacterial and viral infections and aseptic inflammation (Machado et al., 2020; Costa et al., 2022; Senchenkova et al., 2019 Galvão et al., 2017). Recently, it has been demonstrated that AnxA1 suppresses excessive inflammation without changing in viral titers during Dengue and Chikungunya infections in mice (Costa et al; 2022, de Araújo et al, 2022).

Some of the main mechanisms by which AnxA1 curtails inflammation involve modulating neutrophil activity. Seminal works in the field demonstrated that AnxA1 promotes L-selectin shedding in neutrophils, decreases the influx of neutrophils into inflamed tissues, and also stimulates neutrophil apoptosis and efferocytosis (Perretti et al., 1996, Solito et al., 2003, Vago et al., 2012). Neutrophils were found to be crucial cells in COVID-19 pathogenesis (Chen et al., 2020). In addition, rising levels of systemic AnxA1 are correlated with the severity of disease and with high neutrophil counts during COVID-19 (Busch et al., 2022). However, to date, whether and how AnxA1 may operate in the lungs during coronavirus infection and its potential as a therapy against SARS-CoV-2 remain to be determined.

Here, using genetically ablated mice for AnxA1 and different coronaviruses, we demonstrated that endogenous AnxA1 expression increases during the course of infection and that the molecule undergoes cleavage during this process. Our findings revealed that AnxA1 limits neutrophil influx and lung inflammation without impairing the host’s ability to control viral replication. Notably, treatment with the AnxA1-mimetic peptide Ac_2-26_, either alone or in combination with the antiviral drug Remdesivir, significantly reduced the inflammation and lethality caused by SARS-CoV-2 infection in mice.

## Results

### 1. AnxA1 levels increase in conjunction with lung lesions caused by MHV-3

To explore the role of AnxA1 during coronavirus infection, we used a mouse model of intranasal infection with murine betacoronavirus MHV-3 (hepatitis virus 3 strain) (**scheme in Figure 1, A**), in wild-type (WT) mice (Andrade et al., 2021). Given that AnxA1-knockout mice we used are on the BALB/c genetic background (Machado et al, 199), we first characterized the MHV-3 infection in BALB/c mice. Supplementary Figure 1 shows the characterization of the intranasal infection with three different inocula (3 × 10^1^, 3 × 10^2^ and 3 × 10^3^ PFU/mouse) of MHV-3. The intermediate inoculum (3×10^2^ PFU/mouse) was selected for its optimal time window for assessing disease parameters. An inoculum of 3×10^2^ PFU/mouse of MHV-3 caused body weight loss, lung lesion, production of inflammatory mediators in the lungs (TNF, IL-6 and CXCL-1), viral detection by qPCR assay and death of the animals 5 - 6 days post infection (**Supplementary Figure 1, A - G**).

**Figure 1:**
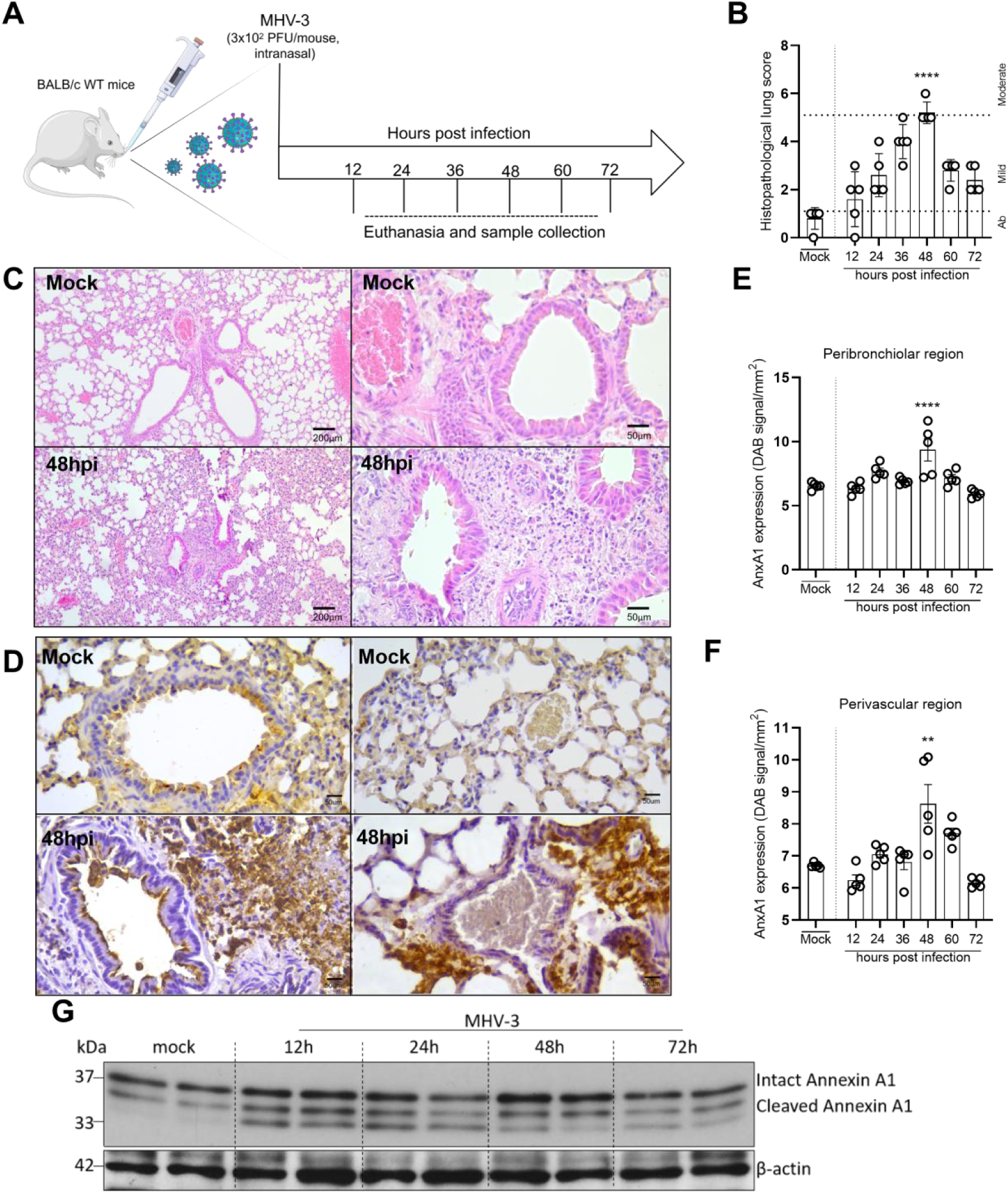
AnxA1 expression is increased during lung inflammation provoked by murine coronavirus (MHV-3) in mice. (A) Experimental design. (B) Histopathological scores of lung sections stained with Hematoxylin and Eosin (n=5). In the graph, “Ab” means absence of lung lesions. (C) Representative images of lung sections (mock and 48hpi group) stained by Hematoxylin and Eosin (Scale Bar 200 μm and 50 µm). (E and F) AnxA1 expression evaluated by immunohistochemistry in the peribronchiolar region and perivascular region. (D) Representative images of lung sections stained by immunohistochemistry to detect AnxA1 in BALB/c wild type (WT) mice (n=5) (Scale Bar, 50 μm). (G) Representative Western blot for AnxA1 at different time points post infection (n=2). ****:p<0.0001 (B and E, mock vs. 48hpi); **:p<0.01 (F, mock vs. 48hpi). Results are shown as mean ± SEM. In (B), data were analyzed by Kruskal-Wallys followed by Dunn’s multiple comparison test. In (E), One Way ANOVA followed by Holm-Sidak’s multiple comparison tests were employed. In F, One Way ANOVA followed by Sidak’s multiple comparison test were employed.

Histopathological analysis showed lung lesions characterized by peribronchiolar and perivascular leukocyte infiltration, peaking at 48 hours post-infection (48 hpi) (**Supplementary Figure 1, C**), (**Figure 1, B**). Immunohistochemistry assays showed increased global expression of AnxA1, particularly in the peribronchiolar and perivascular areas at 48 hpi, which coincided with the maximum histopathological inflammatory scores and leukocyte infiltration (**Figure 1, B and C**). Western blot analysis further detected both full-length AnxA1 and its cleavage fragments in lung tissue extracts, corroborating the immunohistochemical findings (**Figure 1, G**).

### 2. Enhanced inflammatory responses in AnxA1-deficient mice infected with MHV-3

Given the observed increase in AnxA1 levels in the lungs of MHV-3 infected mice and its known role in contain exacerbated inflammation in different contexts, we evaluated the possible contribution of this protein during MHV-3 infection in AnxA1 genetic ablated mice (AnxA1KO) (**scheme in Figure 2, A**). Interestingly, virus replication was similar in the lungs of wild-type (WT) and AnxA1KO mice (**Figure 2, B).** Despite this, AnxA1KO mice showed increased histopathological scores and vascular leakage at the peak of the lung lesion (48 hpi) (**Figure 2 C, D and E)**. Notably, all AnxA1KO mice presented moderate lung lesions, with scores exceeding 5, compared to the mild lesions observed in WT mice (**Figure 2D**). No differences between WT and AnxA1KO groups were observed regarding TNF and IL-6 levels in the lungs, while CXCL1 levels were significantly increased in the absence of AnxA1 in comparison to infected WT littermates (**Figure 2 F, G and H**). Overall, these findings demonstrate that AnxA1 is not essential to control of MHV-3 replication but plays a key role in modulating leukocyte accumulation in the lungs.

**Figure 2:**
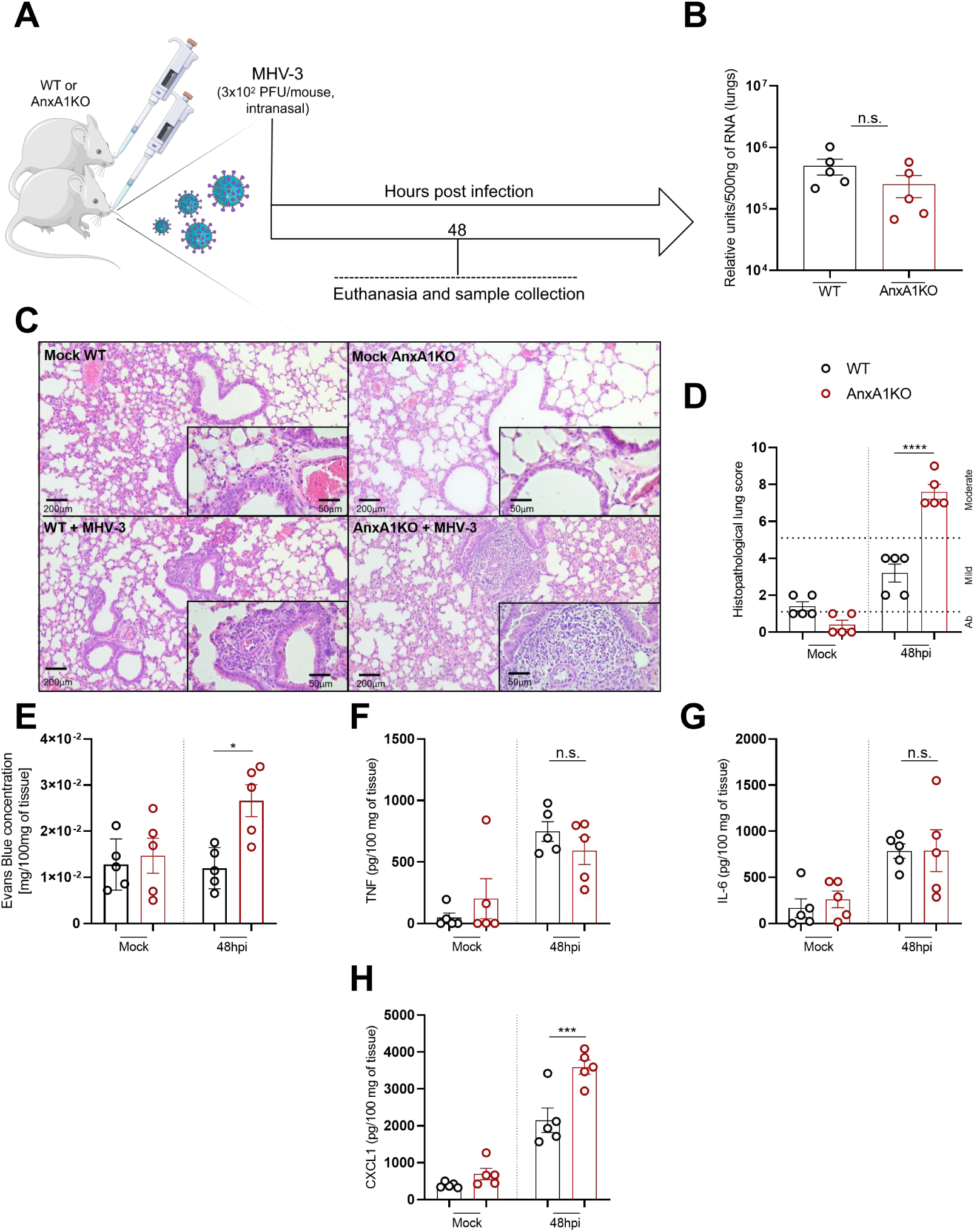
Deficiency of AnxA1 exacerbates lung inflammation without interfering with viral replication. (A) Experimental design. (B) Viral detection by qPCR assay 48 hours post infection. Results are expressed as relative units per 500 ng of RNA used to produce cDNA and are representative of two different experiments (n=5). (C) Representative images of lung sections stained by Hematoxylin and Eosin (Mock group and 48 hours post infection (48hpi)) (Scale Bar 200 μm and 50 μm).(D) Histopathological scores in lung sections (n=5) of BALB/c WT and AnxA1KO mice. (E) Vascular leakage evaluated by Evans Blue concentration assay in the lungs. Concentration of TNF, IL-6 and CXCL1 in the lungs (F, G and H respectively) (expressed as pg/mL of homogenized tissue supernatant) (n=5). (H) ***: p<0.01 (WT vs. AnxA1KO 48hpi). **:p<0.0001 (WT vs. AnxA1KO 48hpi in B and G). n.s.: not significant. In (B), Mann-Whitney test was used to assess the differences between the groups. From (D) to (H), Two-Way ANOVA tests followed by Tukey’s multiple comparison test were used to assess the differences between the groups. Results are expressed as mean ± SEM.

Given the increased levels of CXCL1 in the lungs of AnxA1KO mice and the well-established role of neutrophils in the context of coronavirus infection, we sought to quantify neutrophils in the lungs of mice during the peak of inflammation, by immunohistochemistry analysis (**Figure 3, A**). Higher numbers of Ly6G+ cells were found in the lungs of AnxA1KO mice (**Figure 3, B**), particularly in lung regions commonly affected by MHV-3 infection, as evidenced by the previous results regarding histopathological analyses: peribronchiolar and perivascular region (**Figure 3, C and D**).

**Figure 3:**
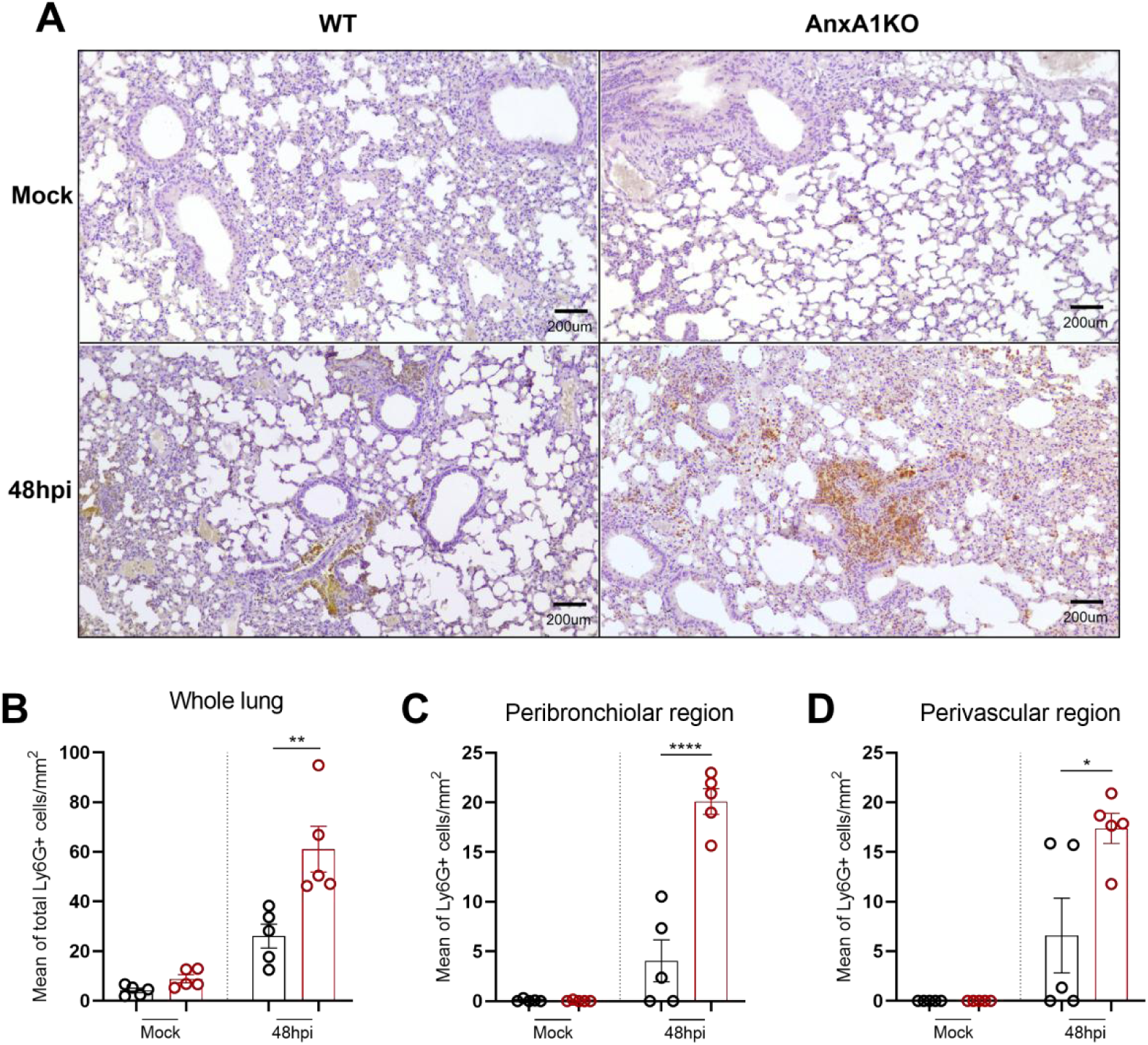
Endogenous AnxA1 limits neutrophil influx in the lungs during MHV-3 infection. The number of neutrophils in the lungs were analyzed using the Ly6G marker for murine neutrophils, by immunohistochemistry (A). Lung sections were obtained from WT and AnxA1KO mice in paraffin-embedded tissues (n=5 per group). Total neutrophils (B) are the sum of neutrophils quantified in four main areas of the lungs: alveolar parenchyma, perivascular region, intravascular region and peribronchiolar region. (B) and (C) shows the levels of neutrophils in peribronchiolar and perivascular regions. Positive cells were manually counted and results are expressed by the mean of Ly6G+ cells per mm2 of lung tissue. All data were analyzed by Two-Way ANOVA and Tukey’s multiple comparison test for both lung areas. ****: p<0.0001. **:p=0.0013. *:p=0.01. All comparisons refer to differences between WT and AnxA1KO mice at 48hpi. Results are expressed as mean ± SEM.

### 3. SARS-CoV-2 infection is associated with increased cleavage of AnxA1 in K18-hACE2 mice

The MHV-3 infection model effectively replicates various COVID-19 parameters in mice, offering valuable insights (Andrade et al., 2021). However, fundamental differences exist between MHV-3 and SARS-CoV-2, especially the mechanisms of viral entry in host cells (Hirai et al., 2010; Zhou et al., 2020). Based on our previous findings on AnxA1 expression during MHV-3 infection, we investigated whether SARS-CoV-2 infection similarly affects AnxA1 expression or its cleavage in the lungs of mice. To this end, K18-hACE2 mice, which overexpress human ACE2 in epithelia cells and are susceptible to SARS-CoV-2 infection, were infected with 3×10⁴ PFU/mouse as previously described (Lima et al., 2024) (illustrated in **Figure 4, A**). In this model, the peak of viral detection occurs 3 days post-infection (dpi), while the maximum lung lesion scores are observed at 5 dpi. Unlike the MHV-3 model, which exhibits intense inflammation in peribronchiolar and perivascular regions, SARS-CoV-2 infection in K18-hACE2 mice triggers a more diffuse pattern of leukocyte infiltration across the alveolar parenchyma. To assess AnxA1 expression under these conditions, we conducted immunohistochemical analyses focused on these regions. The results showed a significant increase in AnxA1 expression 5 days post-infection (**Figure 4, B**). Western blot analysis of lung whole protein extracts further revealed that AnxA1 is cleaved overtime following SARS-CoV-2 infection (**Figure 4, C-F**), in a time point associated with higher inflammatory response (Lima et al, 2024). This observation was reinforced by measuring the ratio between cleaved to full-length AnxA1 (**Figure 4, E**). In summary, these findings suggest that AnxA1 cleavage is associated with lung inflammation in SARS-CoV-2-infected K18-hACE2 mice, mainly at the peak of leukocyte infiltration and lung lesions (Lima et al., 2024).

**Figure 4:**
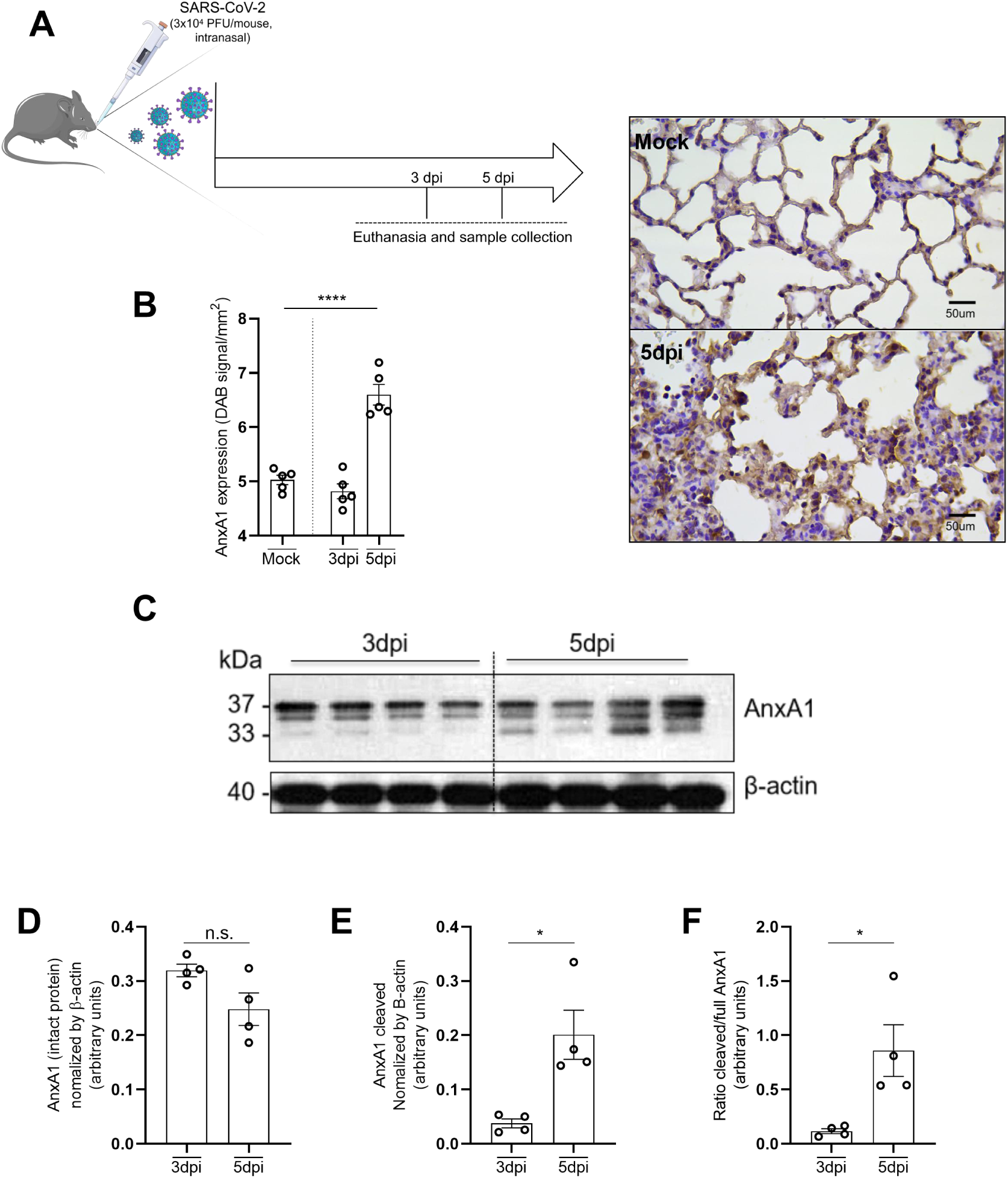
AnxA1 expression during SARS-CoV-2 infection and pattern of cleavage of AnxA1 in in the lungs of K18-hACE2 mice. (A) Experimental design. K18-hACE2 mice (n=5) were intranasally infected with 3×10^4^ PFU/mouse of SARS-CoV-2 gamma variant and euthanized at 3 and 5 days post infection (dpi). (B, left) AnxA1 expression was evaluated by immunohistochemistry in the lungs of K18-hACE2 mice (n=5). (B, right) Representative images of lung sections immunostained with AnxA1 antibody (400x magnification, Scale Bar, 50 μm). (C) Representative image of the Western blot for AnxA1 at different time points post infection in the lungs of SARS-CoV-2-infected mice. In (D), quantification of intact AnxA1. In (E), quantification of cleaved AnxA1. In (F), ratio between cleaved and full AnxA1. In (B), ****p<0.0001 between Mock and 5dpi groups. In (E), *p=0.01 between 3dpi and 5dpi groups; In (F), *p=0.02 between 3dpi and 5dpi groups. In (B), One-Way ANOVA followed by Sidak’s multiple comparison tests were used to evaluate the differences between the groups.In (D-F), Unpaired t test was used to evaluate the differences between the groups. Results are expressed as mean ± SEM.

### 4. Treatment with the AnxA1-mimetic peptide Ac_2-26_ decreased lung lesion and rescued mice from lethality caused by SARS-CoV-2 infection

Ac_2-26_ is a mimetic peptide derived from amino-terminal portion of AnxA1 shown to recapitulate a sort of pro-resolving/anti-inflammatory effects of AnxA1 (Costa et al., 2022; de Araújo et al., 2022; Sugimoto et al., 2016). Based on our previous results showing the importance of AnxA1 in suppress excessive inflammation during MHV-3 infection and the increased cleavage of AnxA1 in the lungs of SARS-CoV-2 infected mice, we explored whether AnxA1-mimetic peptide Ac_2-26_ could be beneficial against SARS-CoV-2 infection. Due to the importance of SARS-CoV-2 to human disease, we tested the potential of Ac_2-26_ in K18-hACE2 transgenic mice that were infected with the gamma variant of SARS-CoV-2 (3×10^4^PFU/mouse). Mice were treated with Ac_2-26_ intraperitoneally every 12h, starting 36 hours post infection (**scheme in Figure 4, A**). Akin to what was observed for AnxA1KO and WT mice, no differences were found between infected groups concerning viral detection assessed by qPCR assay in the lungs (**Figure 4, B**). Ac_2-26_ treatment of infected mice promoted a decreased lung inflammation/leukocyte infiltrate induced by SARS-CoV-2 (**Figure 4, C and D**). The treatment significantly decreased the clinical score (disease signs) and lethality associated with SARS-CoV-2 infection (30 % of survival in the Ac_2-26_ treated group *versus* 10% of survival in the vehicle treated group) (**Figure 4, D and E, respectively**). These results indicate that Ac_2-26_ is able to modulate cell infiltration and curtail lung inflammation without affecting the host’s ability to manage viral replication in the lungs. Additionally, Ac_2-26_ affords significant protection against lethality induced by SARS-CoV-2.

### 5. Ac_2-26_ and Remdesivir provide similar protection against inflammatory lung infiltrates and lethality caused by SARS-CoV-2 infection

The development of host-targeted therapies against infectious diseases are critical for human health (Tavares et al, 2022). Herein, we specifically compared the efficacy of Ac_2-26_ to one of the most common antivirals against SARS-CoV-2, the nucleotide analogue Remdesivir. Additionally, we investigated whether combining Ac_2-26_ with Remdesivir could enhance protection and mitigate disease severity associated with SARS-CoV-2 infection in K18-hACE2 mice (**Scheme in Figure 5, A**). At 3 dpi, Remdesivir treatment significantly reduced SARS-CoV-2 replication in the lungs, which was not observed in the group treated with Ac_2-26_ alone (**Figure 5, B**). In addition, at 3dpi Remdesivir either alone or in combination with Ac_2-26_ was able to decrease lung inflammation (**Figure 5, C and D**). At 5 dpi, all treatments were able to decrease inflammation when compared to the vehicle treated group (**Figure 5, E**). Although the evaluation of the clinical score showed no statistical difference among the treated groups (**Figure 5, F**), the association of Remdesivir to Ac_2-26_ rescued 50% of mice from lethality provoked by SARS-CoV-2 infection (**Figure 5, G**). Remdesivir and Ac_2-26_ alone were able to rescue 40% and 30% of mice, respectively, from SARS-CoV-2-indecd lethality. No statistically significant differences were found among all treated groups. Altogether, these findings demonstrate that Ac_2-26_ and Remdesivir exert similar protection against lung inflammation and lethality induced by SARS-CoV-2. Combining Remdesivir with Ac_2-26_ treatment did not afford further protection than using either treatment alone. Indeed, combined treatment improved lung inflammation at the peak of lung lesions and lethality to similar levels to those found in either treatments alone.

**Figure 5:**
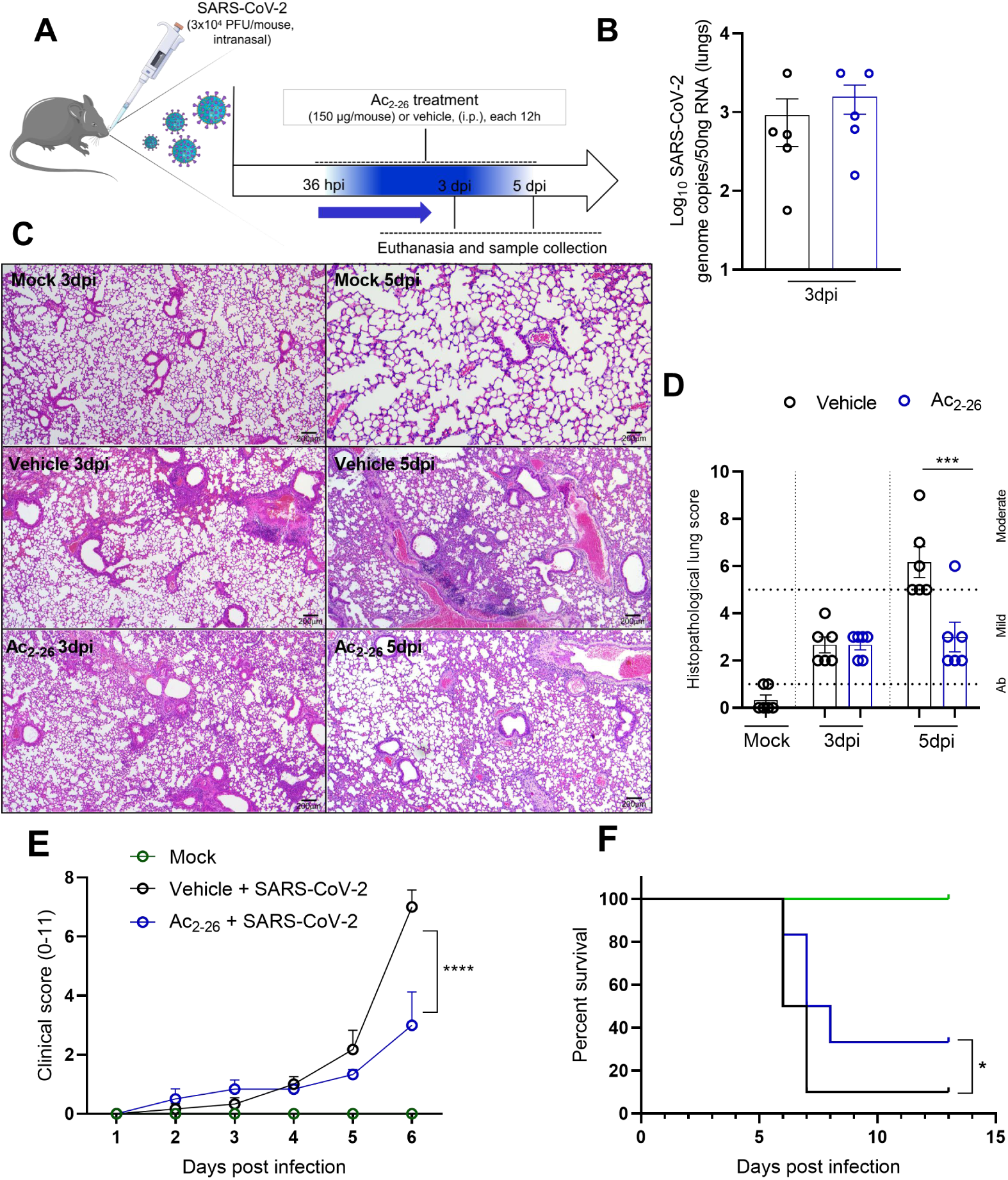
Treatment with the AnxA1 mimetic peptide Ac_2-26_ ameliorates lung damage and clinical score protecting mice from lethality induced by SARS-CoV-2 infection in k18-hACE2 mice. (A) Experimental design. k18-hACE2 transgenic mice (n=6) were intranasally infected with 3×10^4^ PFU/mouse of SARS-CoV-2 and treated with Ac_2-26_ or vehicle starting 36 hours post-infection (36 hpi) and given every 12 hours until euthanasia (3 dpi and 5dpi). (B) Ac_2-26_ does not alter the ability of the host to deal with viral replication in the lungs, as no differences were observed between Ac_2-26_ treated or vehicle treated groups by qPCR assay. (C and D) Ac_2-26_ treatment decreased inflammatory infiltration provoked by SARS-CoV-2 in k18-hACE2 mice at 5dpi (n=6). Ac_2-26_ treatment decreased clinical score and lethality provoked by SARS-CoV-2 (n=5-9) (E and F, respectively). Differences between the groups in (D) were evaluated by Two-Way ANOVA followed by Tukey’s multiple comparison test (***p=0.033 Vehicle vs. treated group at 6dpi). In (E), differences were evaluated using Two-Way ANOVA followed by Tukey’s multiple comparison test (***p<0.0001; Vehicle vs Ac_2-26_ treated groups at 5dpi). In (F) survival curves of infected mice treated or not with Ac_2-26_ (Kaplan-Meyer survival plot), the differences were evaluated by using Gehan-Breslow-Wilcoxon test (*p=0.04). Results are expressed as Mean ± SEM.

**Figure 6:**
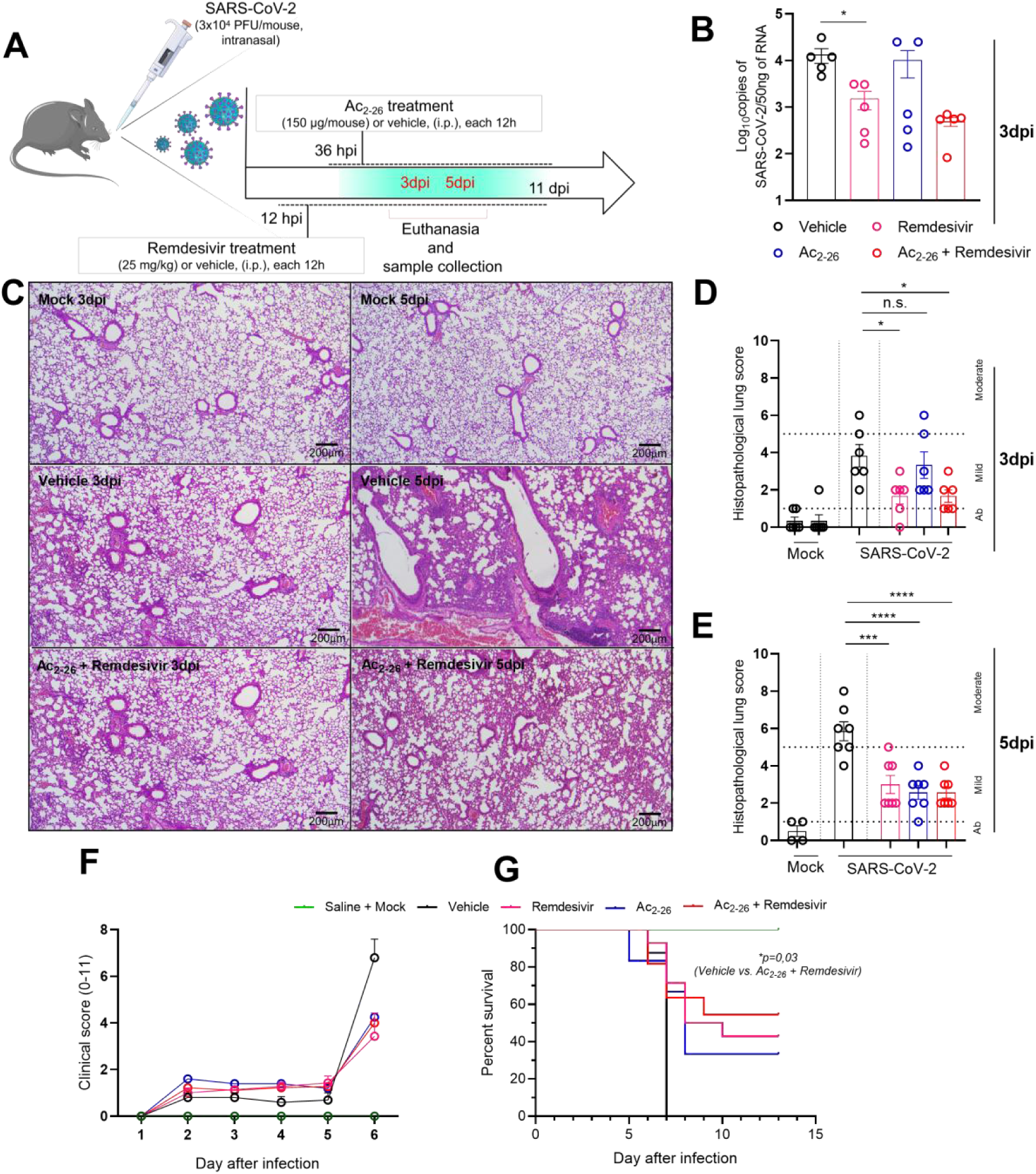
The addition of Remdesivir to Ac_2-26_ protects mice from lung inflammation and lethality induced by SARS-CoV-2 infection. (A) Experimental design. Mice were infected with 3×10^4^ PFU/mouse and treated with Remdesivir (starting at 12 hpi, each 12h) and Ac_2-26_ (starting at 36hpi, each 12h). For evaluation of infection-induced lethality, animals were treated until 11 dpi. (B) Viral detection in the lungs was assessed by qPCR assay 3 days after infection. (D-F) Lung sections of infected mice were evaluated for the histopathological scores at 3 and 5 dpi. Remdesivir + Ac_2-26_ treatment reduced 50% of lethality evoked by SARS-CoV-2 infection (G). In (B), *p=0.02 between Vehicle and Remdesivir treatment groups. In (D),*p=0,01 between Vehicle and Remdesivir or Vehicle and Ac_2-26_ + Remdesivir. In (E), ****p<0.0001. In (G), *p=0.03 between Vehicle and Ac_2-26_ + Remdesivir groups. In (B), One-Way ANOVA followed by Dunn’s multiple comparison test were used to assess the differences between the groups. In (D), Two-Way ANOVA followed by Tukey’s multiple comparison test were used to assess the differences between the groups. In (E), One-Way ANOVA followed by Sidak’s multiple comparison test were used. In (G), Log-Rank (Mantel-Cox) test was used to assess the differences between the groups. Results are expressed as mean ± SEM.

## Discussion

The complex interplay between pro-resolving mediators and infectious diseases has long been a subject of interest, further emphasized during the emergence of COVID-19 (Russell and Schwarze, 2014; Ferri et al., 2023; Sousa et al., 2020; Costa et al., 2024). Of note, AnxA1 is one of the most studied pro-resolving mediator and holds significant impact in the course of infections (Vago et al, 2021; Tavares et al, 2022). For instance, in preclinical models of infectious deseases, AnxA1 limits excessive inflammation during bacterial and viral infections (Machado et al., 2020; Costa et al., 2022; Schloer et al, 2019; Ni et al., 2021). In this study, we have shown that endogenous AnxA1 protects mice from excessive neutrophilic infiltration and lung inflammation/damage during infection with a murine betacoronavirus. In SARS-CoV-2 infection mouse model, the cleavage and consequent inactivation of AnxA1 is associated with lung inflammation. In addition, the administration of its mimetic peptide, Ac_2-26_, protected mice from lung inflammation and lethality evoked by SARS-CoV-2. Of note, Ac_2-26_ reached similar levels of protection when compared to Remdesivir treatment against SARS-CoV-2 infection.

Different studies explored AnxA1 circulating levels or its mRNA expression in immune cells by using sc-RNA-seq in COVID-19 patients. Bush and colleagues demonstrated increased AnxA1 plasma levels in moderate and severe COVID-19 patients accompanied of higher numbers of circulatory neutrophils (Busch et al., 2022). However, Shenoy and co-workers showed a significant decrease in AnxA1 serum levels in severe COVID-19 patients (Shenoy et al., 2023). Despite both studies declaring that COVID-19 severity was classified by the same criteria (according to the World Health Organization (WHO)), the contrasting results might be explained by a variety of reasons, such as small sample size, different genotypes of the population studied and plasma/serum preference. In the lungs of COVID-19 patients, AnxA1/FPR1 pair interaction was shown to be enhanced during infection and crucial for the interplay between squamous epithelial cells and myeloid cells, such as neutrophils and macrophages (Ren et al., 2021; Qin et al., 2023). However, to date, no study has comprehensively show AnxA1 protein expression in the lungs and its potential against coronavirus infection, either in human autopsy samples or using preclinical models. Our results demonstrate that coronavirus infection increases AnxA1 expression in the lungs at the protein level, especially in areas associated with leukocyte influx, such as peribronchiolar and perivascular regions. These findings might be attributed to an increased influx of leukocytes into the lungs, which express AnxA1, alongside elevated expression by resident cells. In keeping with that, it was demonstrated increased mRNA expression of AnxA1 in squamous epithelial cells during SARS-CoV-2 infection by transcriptomic analysis (Ren et al., 2021).

AnxA1 can be cleaved by different proteases in sites of inflammation, such as Proteinase 3 and neutrophil elastase (Vago et al., 2016; Sugimoto et al., 2016). Evidence suggests that cleaved forms of AnxA1 might sustain pro-inflammatory signaling in different contexts. Examples include LPS-induced pleurisy in mice, neuroinflammation in the neocortex of neurodegenerative patients, cystic fibrosis patients and in BAL fluid of bronchial carcinoma and resolving pneumonia in humans (Vago et al., 2012; Chua et al., 2022; Tsao et al., 2012; Smith et al., 1990). Our findings reveal that SARS-CoV-2 infection leads to AnxA1 cleavage in the lungs with consequent reduction of intact protein of 37 kDa. Noteworthy, AnxA1 cleavage coincides with the peak of lung lesions, suggesting a potential association between the cleaved forms of AnxA1 and excessive inflammation during SARS-CoV-2 infection in K18-hACE2 mice. The significance of AnxA1 cleavage is further highlighted by efforts in the field to develop cleavage-resistant AnxA1 variants aimed at mitigating excessive inflammation (Pederzoli-Ribeil et al., 2010; Dalli et al., 2013).

Seminal studies showed the importance of endogenous AnxA1 levels in restrain leukocyte influx and excessive inflammation into the tissues in different contexts (Guido et al., 2013; Hayhoe et al., 2006; Buss et al., 2015; Walther et al., 2000). Mechanistically, evidence suggests that AnxA1 acts by shedding L-selectin on the surface of leukocytes (Coupade et al., 2003; Solito et al., 2003). In line with previous studies, our results demonstrate that the lack of endogenous AnxA1 in knockout mice leads to increased inflammation, by enhancing vascular leakage, CXCL1 levels and neutrophil influx into the lungs during murine coronavirus infection. This corroborates the recent study by Gong and colleagues, who used MHV-1 strain of betacoronavirus to show that vascular leakage during infection is primarily driven by neutrophils in the lungs (Gong et al., 2023). Conversely, Ac_2-26_ treatment, either alone or in combination with Remdesivir, mitigates exaggerated inflammation in SARS-CoV-2 infected mice, providing protection against infection-associated lethality.

Notably, both endogenous AnxA1 or exogenous Ac_2-26_ administration demonstrated the ability to suppress excessive inflammation without compromising the host’s capacity to control viral replication. This was evident as no differences in viral detection by qPCR were observed between WT and AnxA1KO mice, or between Ac_2-26_ treated group and vehicle group. Developing novel host-targeted therapies for infectious diseases requires a therapeutic candidate capable of striking a precise balance between mitigating excessive inflammatory responses and maintaining the immune system’s effectiveness in controlling pathogen replication. Our group has previously demonstrated this ability of AnxA1 in studies addressing Dengue and Chikungunya viruses as well (Costa et al., 2022; de Araújo et al., 2022). Here, we aimed to enhance the value of this therapeutic strategy by combining Ac_2-26_ treatment with a nucleoside analog. The same approach has been tested in different infectious models with other pro-resolving molecules, such as the combination of Lipoxin A4 or Angiotensin-(1-7) with antibiotics during *Pseudomonas aeruginosa* infection, which increased the capacity of ciprofloxacin and imipenem to kill bacteria (Thornton et al., 2021; Zaidan et al., 2024).

Remdesivir, either alone or combined with Ac_2-26_, prevented lung inflammation at early time points (3 dpi), likely due to its ability to decrease viral loads and consequently the presence of viral PAMPs. The addition of Ac_2-26_ to Remdesivir did not significantly improve protection against SARS-CoV-2-induced lethality. Yet, Ac_2-26_ alone successfully mitigated exaggerated inflammation at the peak of the response (5 dpi), preventing lung lesions without affecting viral detection in the lungs. This demonstrates that Ac_2-26_ exhibits similar levels of protection when compared to one of the most used antivirals against SARS-CoV-2. Alternatively to Remdesivir, Ac_2-26_ is oriented to mitigate excessive inflammation rather than viral replication. This underscores the potential of host-oriented therapies against viral diseases and the importance of containing excessive/misplaced inflammation.

Nonetheless, despite its promising therapeutic potential and possible future applications, caution is essential when using pro-resolving drugs to treat infectious diseases. For example, Lipoxin A4 treatment during the early stages of pneumosepsis reduced cell migration and exacerbated the infection. In contrast, when administered at later stages post-infection, Lipoxin A4 improved survival rate by mitigating the excessive inflammatory response (Sordi et al., 2013). This was also verified during IAV infection in mice, in which a stable LXA4 analogue decreased leukocyte infiltration and inflammation in the lungs (Rago et al., 2024, preprint). Here, we opted to initiate Ac_2-26_ treatment 36 hours post-infection (1.5 dpi), as the clinical score of infected K18-hACE2 mice begins to slightly increase by day 2 post-infection. Our group has previously demonstrated that early treatment with immunomodulatory drugs against coronavirus infection, such as the PDE4 inhibitor Roflumilast, can exacerbate the infection (Beltrami et al., 2024). Therefore, the effectiveness of immunomodulatory drugs against infectious agents is fundamentally time-dependent.

It must be noted that, despite its encouraging prospects, the role of AnxA1 can vary depending on the pathogen. For instance, both HSV-1 and H1N1 exploit the AnxA1 pathway to enter cells via the FPR2 receptor (Wang et al., 2022; Arora et al., 2016). Notably, this does not appear to be the case for coronavirus infections. In murine betacoronavirus infection, AnxA1KO mice showed no differences in viral detection in the lungs when compared to WT counterparts; and Ac_2-26_ treatment during SARS-CoV-2 infection had no impact on viral replication as well.

Here, we demonstrate that the use of a pro-resolving drug exhibit similar protection to pathogen replication inhibitors such as nucleoside analogs, which can positively influence the disease course by controlling misguided inflammation caused by infectious agents. In recent years, various pathogens have emerged and re-emerged, triggering epidemics and creating social and health crises that overwhelm healthcare systems. Developing novel therapies to address dysregulated inflammation caused by pathogens has become imperative. Our findings emphasize the critical role of endogenous AnxA1 in controlling neutrophil influx into the lungs during murine coronavirus infection. Additionally, we highlight its potential as a therapeutic strategy against SARS-CoV-2, paving a way for the use of pro-resolving, host-targeted therapies to mitigate misplaced inflammation caused by novel and recurring viral agents.

### Limitations

As limitations of our work, we acknowledge that MHV-3 is not a natural respiratory viral infection in its natural hosts, as it induces pulmonary disease only when administered intranasally. Nonetheless, various independent groups have successfully employed different strains of MHV-3 to model COVID-like disease in mice for pathogenesis studies and drug tests (Andrade et al, 2021; Yang et al., 2014; Gong et al., 2023; Campolina-Silva et al., 2023). Nevertheless, K18-hACE2 mice overexpress human ACE2 exclusively in epithelial lineage cells, preventing leukocytes and other cell types from being infected by SARS-CoV-2. This limitation means that the model cannot fully replicate the pathogenesis of COVID-19 as it occurs in humans.

### Clinical perspectives

- AnxA1 is a key modulator of the resolution of inflammation, including in the context of infectious models of diseases. However, to date, no study has evaluated the endogenous role of AnxA1 or the therapeutic potential of its mimetic peptide, Ac_2-26_, against coronavirus infection.
- Mice lacking AnxA1 showed increased neutrophil influx and lung inflammation in MHV-3-infected mice, and the exogenous administration of Ac_2-26_ decreased lung inflammation and prevented lethality provoked by SARS-CoV-2 in K18-hACE2 mice.
- Our findings demonstrate that Ac_2-26_ is a potential candidate to target excessive inflammatory responses during SARS-CoV-2 infection, and support further clinical studies to explore its potential as adjuvant treatment in human infections.

### Data availability

The data that support the findings are available from the corresponding author upon reasonable request.

## Materials and Methods

### Mice

For MHV-3 experiments, male and female BALB/c mice were obtained from the Center of Bioterism of Universidade Federal de Minas Gerais (UFMG), Brazil. AnxA1 knockout mice in BALB/c background (Dufton et al., 2003) were bred and maintained at animal facilities of the Department of Biochemistry and Immunology of UFMG. 9-week-old BALB/c WT and AnxA1KO mice were housed in ventilated cages in specific pathogen-free conditions at a constant temperature of 25°C and 12/12 h light/dark cycle, with free access to food and water. Infections with MHV-3 were performed in Animal Biosafety Level 2 facilities (BSL-2) in the Institute of Biological Sciences. For SARS-CoV-2 experiments, male and female transgenic mice expressing the Human Angiotensin Converting Enzyme (K18-hACE2) in the C57BL/6 background were obtained from Jackson Laboratories and maintained at the vivarium of the Biochemistry and Immunology Department at UFMG. 9-weeks-old K18-hACE2 mice were bred at UFMG animal facilities and infections with SARS-CoV-2 were performed in Animal Biosafety Level 3 facilities (BSL-3) at Institute of Biological Sciences of UFMG. All the experiments were conducted in accordance with the recommendations in the Guide for the Care and Use of Laboratory Animals of the Brazilian National Council of Animal Experimentation (CONCEA). The *in vivo* experimental procedures were approved by the UFMG Ethics Committee for the Use of Animals (process no. 249/2020).

### MHV-3 and SARS-CoV-2 infection

Mice were anesthetized intraperitoneally (ketamine [60 mg/kg] xylazine [5 mg/kg]) and received an intranasal inoculation of 30 μl of sterile saline with MHV-3 (3×10^2^PFU/mouse). The same approach was used for SARS-CoV-2 experiments as previously shown by our group (3×10^4^PFU/mouse, Guimarães et al., 2024). For the Mock control group, animals received sterile saline. For MHV-3 experiments, animals were euthanized for organ collection for analyzes in different time points post infection (every 12h for characterization of the infection model in BALB/c mice, and 48 hours post infection for AnxA1KO mice experiments). For SARS-CoV-2 experiments, animals were euthanized 3 and 5 days after infection.

### Ac_2-26_ and Remdesivir treatment

For Ac_2-26_ treatment, the peptide was formulated with hydroxypropyl-β-cyclodextrin (HP-β-cyclodextrin) and sterile saline solution. For each 5.46mg of HP-β-cyclodextrin, 1mg of Ac_2-26_ is used. Mice received 150μg/animal of Ac_2-26_ starting at 36 hours after infection, twice a day (100μL, i.p.) until euthanasia (Costa et al., 2022; de Araújo et al., 2022). For Remdesivir (Veklury ®) treatment, mice also received 25mg/kg twice a day until euthanasia, but treatment started 12 hours after infection.

### Histopathological analysis

Tissues were harvested and fixed in 10% formalin at pH 7.4 for 48h, followed by dehydration in ethanol and embedding in paraffin. Sections of 5μm of lungs were stained with Hematoxylin and Eosin (H&E). The lung slides were examined for determination of the inflammation-mediated injury using a score system previously used in our group for MHV-3 infections (Andrade et al., 2021; Queiroz-Junior et al., 2023).

### Virus detection by qPCR assay

Briefly, viral RNA was extracted from the tissues using QIAmp® Viral RNA kit following manufacturer’s instructions. Lung tissues were macerated using a lysis buffer of the kit, and eluted RNA was quantified using a spectrophotometer (NanoDrop^TM^, Thermo Scientific). The cDNA was synthesized with 500ng of total RNA using the iScript ^TM^ gDNA Clear cDNA Synthesis Kit (BIO-RAD) following manufacturer’s instructions. The cDNA was diluted 1:10 to be used in the qPCR assay. For MHV-3 detection, Fast SYBR TM Green Master Mix (Applied Biosystems TM) was used for the reaction to quantify the N protein gene of the virus. The following primers were used (5 nM): Forward primer 5′-CAGATCCTTGATGATGGCGTAGT-3′; Reverse primer 5′-AGAGTGTCCTATCCCGACTTTCTC-3′. The standard was obtained by extracting RNA from a known amount of PFU from MHV-3 and results were expressed as arbitrary units/500 ng of RNA. For SARS-CoV-2 detection, reaction was made using 2019-nCoV RUO kit (IDT) for N1 region. Standard was produced from a known number of copies by 2019.nCoV_N Positive Control (IDT). Results were expressed in Relative Units/500 ng cDNA.

### ELISA

Cytokines and chemokines associated with inflammatory response in the lungs were measured using ELISA assay. Briefly, lungs were macerated using TissueLyser LT (Qiagen®) and cytokine extraction buffer (100 mM Tris [pH 7.4], 150 mM NaCl, 1 mM EGTA, 1 mM EDTA, 1% Triton X-100, 0.5% sodium deoxycholate, and 1% protease inhibitor cocktail - Sigma). TNF, IL-6, IL-10, CXCL-1 were measured in tissue homogenates using commercially available Mouse DuoSet ELISA Kits (R&D System). Immunoassays were performed according to the manufacturer’s instructions. Results were determined as pg/100mg of tissue.

### Changes in vascular permeability

The extravasation of Evans Blue dye into the lungs was employed as an index of increased vascular permeability as previously described (Costa et al., 2022; Saria and Lundberg, 1983). The quantity of Evans Blue in the tissue was determined by comparing the extracted absorbance to a standard Evans Blue curve, read at 620 nm using a spectrophotometer plate reader. The results are expressed as the amount of Evans Blue per 100 mg of tissue.

### Immunohistochemistry

Lung slides were deparaffinized, hydrated and endogenous peroxidase blockade was performed using 0.3% of hydrogen peroxide in PBS. For AnxA1 detection, antigen retrieval was achieved using citrate solution (0.01M) pH 6.0. For Ly6G detection, antigen retrieval was made using the EDTA buffer (0.00127M) (pH 9.0). Slides were then incubated with primary antibodies anti-AnxA1 or anti-Ly6G (AnxA1 Invitrogen, Cell Signaling, 1:400; Ly6G - Abcam (EPR22909-135, 1:2000). The sections were then incubated with the secondary biotinylated goat-anti-rabbit antibody and reaction was revealed using the VECTASTAIN Elite ABC HRP Kit and stained with DAB chromogenic solution (Sigma-Aldrich). Counterstaining with hematoxylin was subsequently performed. In negative controls, primary antibodies were omitted. The intensity of AnxA1 expression in the tissues was represented as DAB signal/mm^2^ of tissue area as previously described (Cizkova et al., 2021). For the Ly6G marker, results were represented as the mean of Ly6G+ cells/mm^2^.

### Western Blot

To evaluate AnxA1 expression in the lungs, 10 μg of total protein from lung homogenate supernatant was used. Samples were heated at 100°C for 5 minutes before being loaded onto a 15% (for MHV-3 infection) and 10% (for SARS-CoV-2 infection) Acrylamide/Bisacrylamide gel. Electrophoresis was conducted at 60V for stacking and 100V for 2 hours for resolving steps. Proteins were transferred to a nitrocellulose membrane using 350mA and 140V for 1 hour and 20 minutes. (Transfer buffer Tris 0.025M, Glycine 0.192M, Methanol 4.943M). Membranes were blocked with a solution of PBS with 0.1% Tween 20 and 3% BSA for 1 hour at room temperature. They were then incubated overnight with anti-AnxA1 primary antibody (#71-3400, Invitrogen) at 1:1000 dilution in PBS with 0.1% Tween 20 and 3% BSA. After washing, membranes were incubated with a secondary antibody (Cell Signaling, anti-rabbit IgG) at 1:3000 dilution in the same buffer for 1 hour at room temperature. Immunoreactive bands were detected using an enhanced chemiluminescence (ECL) detection system (GE Healthcare, Piscataway, NJ, USA) (250 μL each of solutions A and B) for 5 minutes at room temperature. Finally, the membrane was exposed to X-ray film in a darkroom for variable times. For densitometric analysis, membranes were evaluated using Fiji imaging software, and normalized by β-actin. Values are expressed as arbitrary units.

### Data and Statistical analysis

GraphPad Prism version 8.0.2 was used for statistical analysis and data plotting. Depending on whether the data distribution was parametric or non-parametric, the following tests were applied: Kruskal-Wallis followed by Dunn’s multiple comparison test, One-Way ANOVA followed by Holm-Sidak and Two-Way ANOVA followed by Tukey’s multiple comparison test. Survival curves were generated using the Kaplan-Meier method, with significance of differences calculated by the Log-Rank (Mantel Cox) or Gehan-Breslow Wilcoxon tests. Results were expressed as mean ± SEM. Differences were considered statistically significant if p<0.05.

## Acknowledgments

We are thank to Ilma Marçal de Souza, Rosemeire Oliveira, Tânia Colina and Letícia Cardoso for their technical assistance. We also thank “Para mulheres na Ciência Prize” provided by L’oréal, UNESCO and ABC for VVC. Thanks are also due to the animal biosafety level 3 laboratory at UFMG (Laboratório Institucional de Pesquisa, LIPq); Centro de Laboratórios Multiusuários, CELAM, Laboratório de Biossegurança Nível 3, NB3-ICB. This work was supported by grants from Coordenação de Aperfeiçoamento de Pessoal de Nível Superior – CAPES/Brazil (Projeto: CAPES - Programa: 9951 - Programa Estratégico Emergencial de Prevenção e Combate a Surtos, Endemias, Epidemias e Pandemias AUX 0641/2020 - Processo 88881.507175/2020-01) and CAPES 11/2020 Epidemias, N° 88887.506690/2020-00. This work was also funded by Instituto Nacional de Ciência e Tecnologia em Dengue e Interação Microrganismo-Hospedeiro for the fundings provided (Processo CNPQ: 465425/2014-3).

---

**Supplementary Figure 1:**
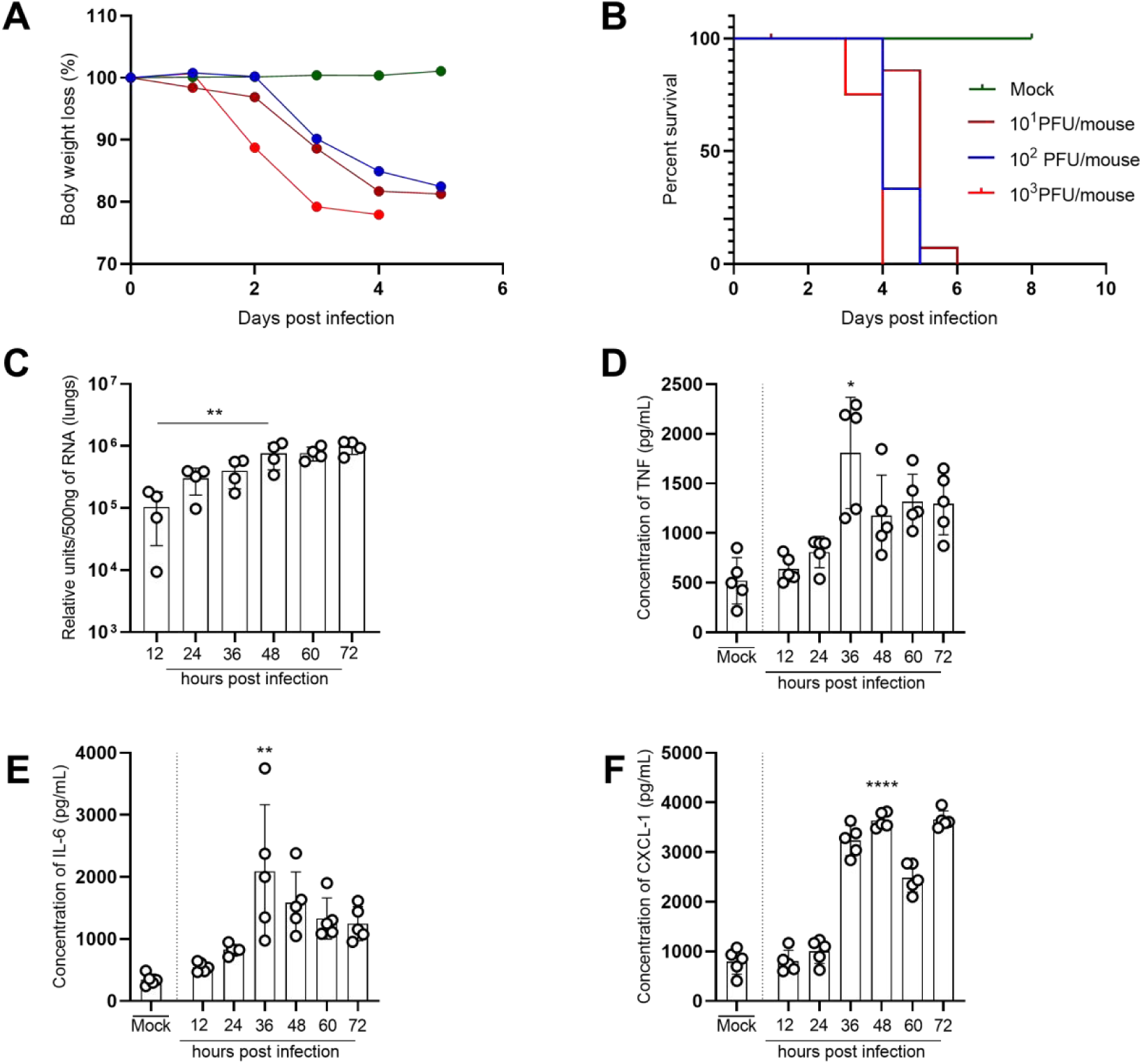
Murine Coronavirus (MHV-3) infection mimics disease aspects of severe COVID-19 in BALB/c mice. (A) Body weight loss and (B) percent survival of BALB/c WT mice infected with different inocula of MHV-3 (3×10^1^ - 3×10^3^PFU/mouse, intranasal). (C) to (F), disease parameters of mice infected with 3×10^2^PFU/mouse. (C) Viral detection by qPCR assay at different time points post infection (results are expressed as relative units per 500ng of RNA used to produce cDNA). Concentration of TNF (D), IL-6 (E) and CXCL-1 (F) (pg/mL) evaluated by ELISA assay in the lungs of mice. *: p=0,0181 (mock vs. 48hpi); **: p=0,0018 (mock vs. 48hpi); ****: p<0,0001 (mock vs. 48hpi). Results are expressed as mean ± SEM.

